## Supplementary material for "High-throughput profiling of sequence recognition by tyrosine kinases and SH2 domains using bacterial peptide display": Figure supplements

### Figure supplements for:

**ORCiDs**

Allyson Li: 0000-0003-2359-7703

Rashmi Voleti: 0000-0002-3705-7460

Minhee Lee: 0000-0002-7141-7351

Dejan Gagoski: 0000-0001-6194-8514

Neel H. Shah: 0000-0002-1186-0626

**A**

| | -5 | -4 | -3 | -2 | -1 | 0 | +1 | +2 | +3 | +4 | +5 |
|---|---|---|---|---|---|---|---|---|---|---|---|
| **Y** | 8779 | 15927 | 5617 | 10733 | 6624 | 800896 | 11063 | 10129 | 11814 | 8603 | 14957 |
| **W** | 33468 | 29232 | 75960 | 53667 | 30871 | 9 | 26426 | 24256 | 23063 | 62469 | 59524 |
| **F** | 40713 | 26039 | 12478 | 17141 | 16764 | 736 | 16012 | 27990 | 30045 | 21315 | 22750 |
| **I** | 29589 | 15068 | 12943 | 18649 | 5553 | 133 | 42501 | 45385 | 13034 | 29318 | 16808 |
| **L** | 125560 | 70574 | 91544 | 93152 | 126228 | 655 | 52276 | 106984 | 100535 | 58098 | 107100 |
| **V** | 162844 | 82964 | 38626 | 24160 | 185975 | 2488 | 65366 | 86110 | 145396 | 35768 | 30867 |
| **M** | 74141 | 21961 | 63128 | 40292 | 22496 | 37 | 48479 | 40730 | 15633 | 34989 | 32642 |
| **P** | 16106 | 49524 | 38437 | 62591 | 28561 | 21 | 22629 | 35166 | 47501 | 25726 | 51929 |
| **G** | 50350 | 89168 | 54565 | 36457 | 68795 | 73 | 90550 | 74862 | 65791 | 76584 | 38873 |
| **A** | 44896 | 79015 | 33562 | 23762 | 82069 | 19 | 32826 | 40414 | 95485 | 24011 | 25759 |
| **T** | 23086 | 37808 | 73126 | 60133 | 17794 | 17 | 50300 | 51455 | 22999 | 45932 | 47245 |
| **S** | 68173 | 83695 | 74004 | 80335 | 56220 | 236 | 67596 | 70852 | 66177 | 101132 | 83396 |
| **C** | 12000 | 13911 | 11670 | 20432 | 7531 | 688 | 16813 | 19391 | 12387 | 39426 | 22015 |
| **N** | 6652 | 12597 | 5638 | 13608 | 2670 | 775 | 24730 | 18664 | 4715 | 11347 | 10767 |
| **Q** | 6585 | 15766 | 15930 | 27609 | 16621 | 4 | 14906 | 10291 | 16292 | 9543 | 34551 |
| **D** | 9917 | 27300 | 4918 | 5877 | 12330 | 2196 | 20848 | 16370 | 23887 | 7257 | 8592 |
| **E** | 27071 | 37753 | 17085 | 11685 | 57762 | 12 | 28582 | 15980 | 48291 | 10442 | 19238 |
| **H** | 2818 | 12539 | 3996 | 12678 | 4241 | 547 | 10843 | 10945 | 10247 | 7129 | 13001 |
| **R** | 37609 | 53006 | 133999 | 148357 | 31100 | 98 | 119354 | 81679 | 33359 | 178733 | 112942 |
| **K** | 14389 | 19925 | 26725 | 32520 | 11306 | 4 | 37224 | 15378 | 7652 | 14504 | 26382 |
| **stop** | 16140 | 17125 | 16946 | 17059 | 19386 | 1253 | 11573 | 7866 | 16594 | 8571 | 31559 |

**B**

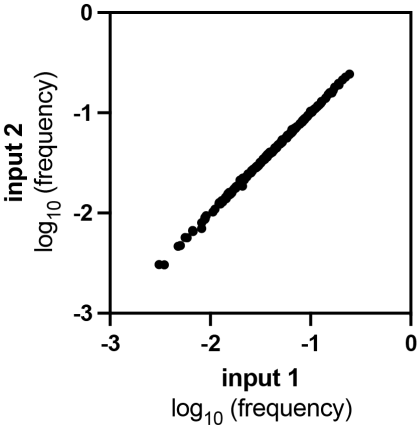


**C**

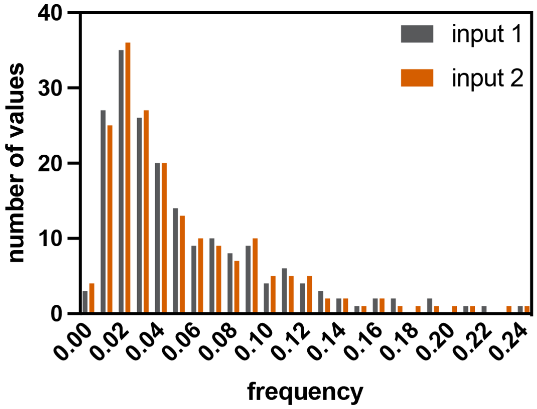


**Figure 1-figure supplement 1. Composition of the $X_5$-Y-$X_5$ library.** (**A**) Table showing the read counts for all amino acids and the stop codon across all positions in the strep-tagged $X_5$-Y-$X_5$ library, from one sequencing run with an unselected (input) library. (**B**) Correlation of amino acid frequencies at each position from two replicates of the input library. (**C**) Distribution of frequencies from two replicates of the input library.

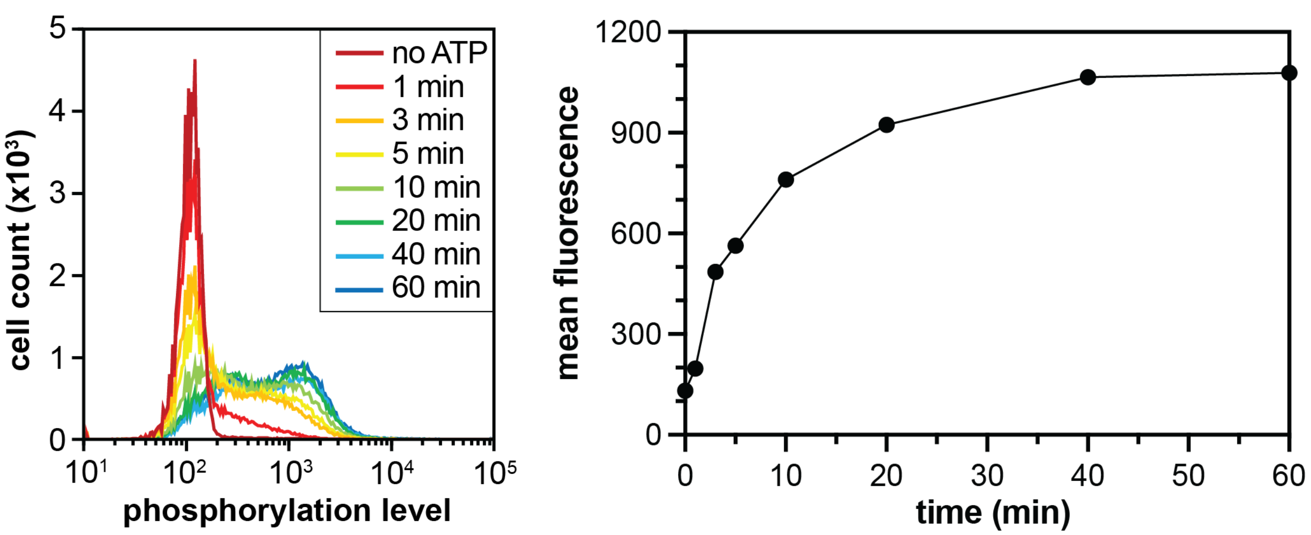


**Figure 1-figure supplement 2. Phosphorylation of the $X_5$-Y-$X_5$ library by c-Src.** Flow cytometry analysis monitoring the distribution of phosphotyrosine levels over time (*left*). The mean fluorescence intensities, which represent phosphorylation levels, plotted as a function of time (*right*).

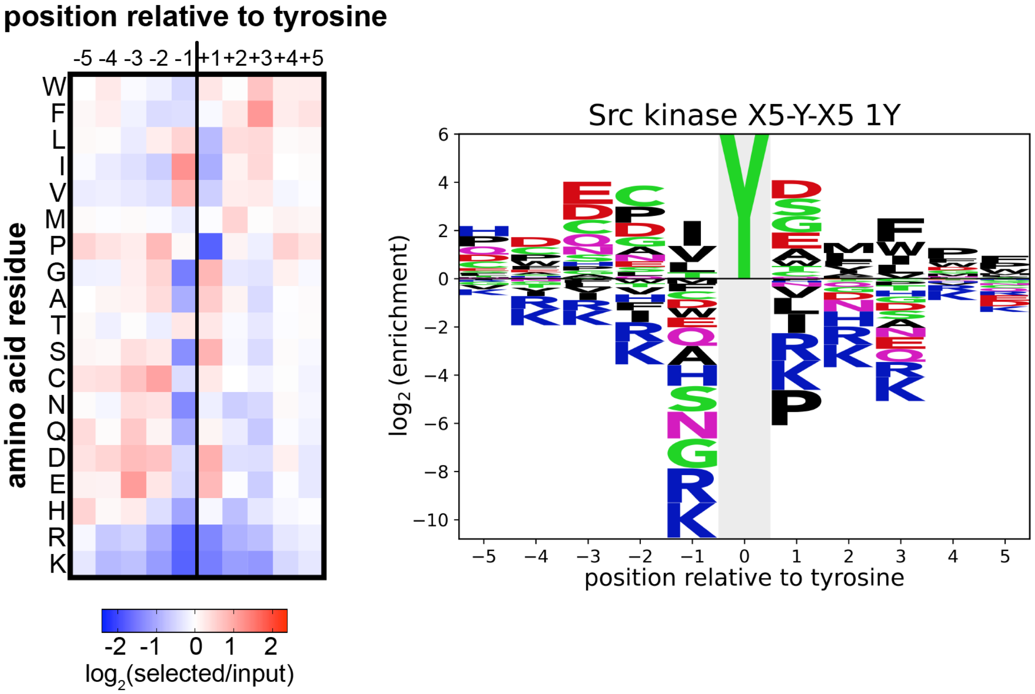


**Figure 1- figure supplement 3. Heatmap and logo depicting the specificity of the c-Src kinase domain, measured using the $X_5$-Y-$X_5$ library.** Only peptides with one central tyrosine were considered in this analysis. Enrichment scores were $log_2$-transformed and are displayed on a color scale from blue (disfavored sequence features, negative value), to white (neutral sequence features, near zero value), to red (favored sequence features, positive value). The same values were used to plot the heatmap and the sequence logo. The height for the central "Y" in the sequence logo is an arbitrary value, chosen for optimal visualization of other features. Values are the average of three replicates.

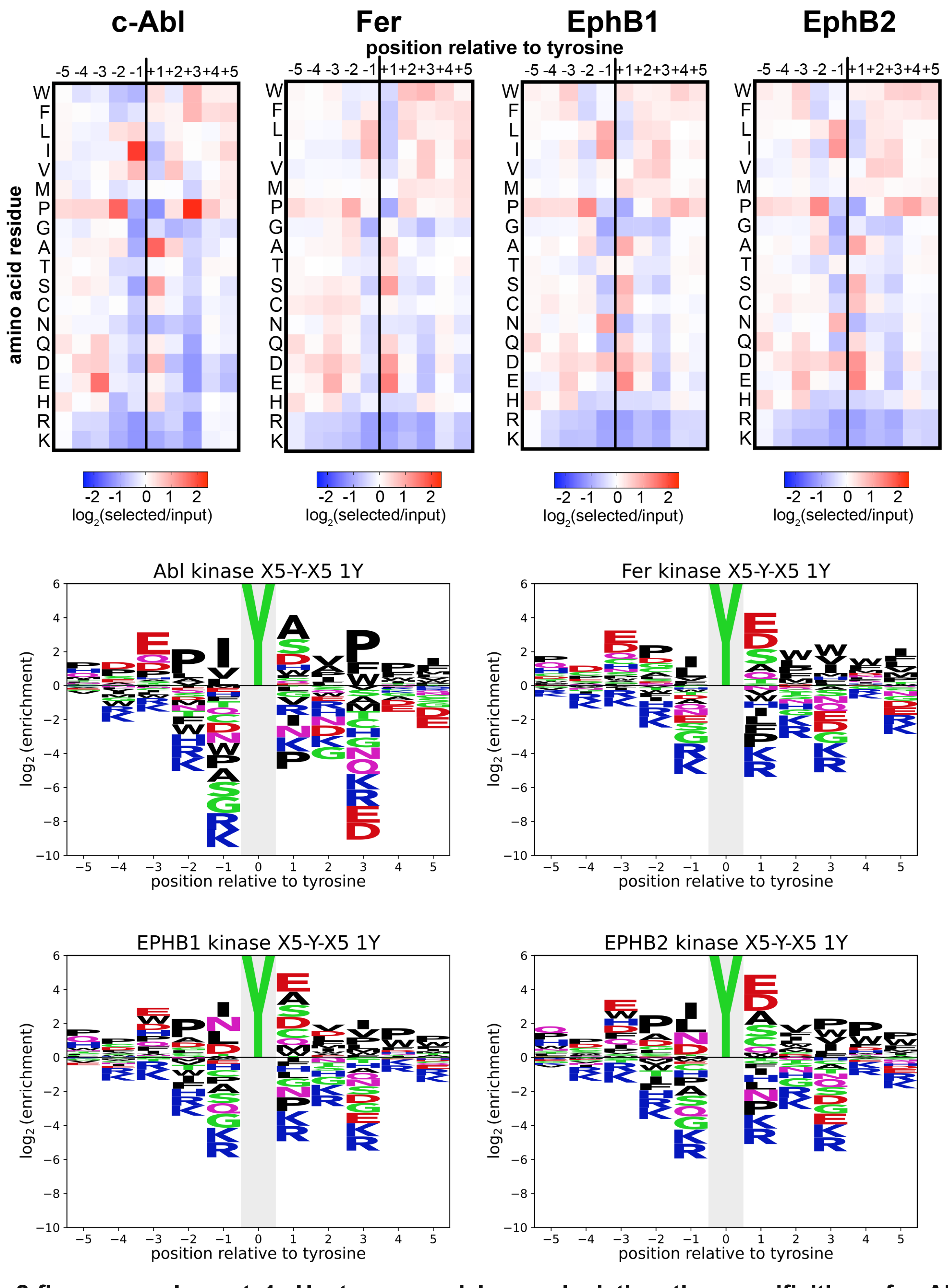


**Figure 2-figure supplement 1. Heatmaps and logos depicting the specificities of c-Abl, Fer, EPHB1, and EPHB2.** Only peptides with one central tyrosine were considered in this analysis. Enrichment scores were $log_2$-transformed and are displayed on a color scale from blue (disfavored sequence features, negative value), to white (neutral sequence features, near zero value), to red (favored sequence features, positive value). The same values were used to plot the heatmaps and the sequence logos. The height for the central “Y” in the sequence logos is an arbitrary value, chosen for optimal visualization of other features. Values are the average of three replicates.

| entry | kinase | peptide name | peptide sequence | $k_{cat}$ ($s^{-1}$) | $K_M$ (μM) |
|---|---|---|---|---|---|
| 1 | c-Src | Src Consensus | GPDECIYDMFPFKKKG | 4.9 ± 0.4 | 196 ± 38 |
| 2 | c-Src | Src Consensus (P-5C, D+1G) | GCDECIYGMFPFKKKG | 4.4 ± 0.2 | 97 ± 10 |
| 3 | c-Src | SrcTide (1995) | GAEEEIYGEFEAKKKG | 3.1 ± 0.2 | 64 ± 10 |
| 4 | c-Src | SrcTide (2014) | GAEEEIYGIFGAKKKG | 1.8 ± 0.1 | 7 ± 3 |
| 5 | c-Src | Fer Consensus | GPDEPIYEWWWIKKKG | 0.4 ± 0.1 | 8 ± 4 |
| 6 | c-Src | Abl Consensus | GPDEPIYAVPPIKKKG | 2.0 ± 0.2 | 159 ± 31 |
| 7 | c-Abl | Abl Consensus | GPDEPIYAVPPIKKKG | 3.0 ± 0.2 | 6 ± 2 |
| 8 | c-Abl | AblTide (2014) | GAPEVIYATPGAKKKG | 2.5 ± 0.2 | 35 ± 8 |

**Figure 2-figure supplement 2. Table of Michaelis-Menten parameters for consensus peptides against c-Src and c-Abl kinase domains.** All measurements were carried out using the ADP-Quest assay in 3-5 replicates. Errors represent the standard error in global fits of all replicates to the Michaelis-Menten equation.

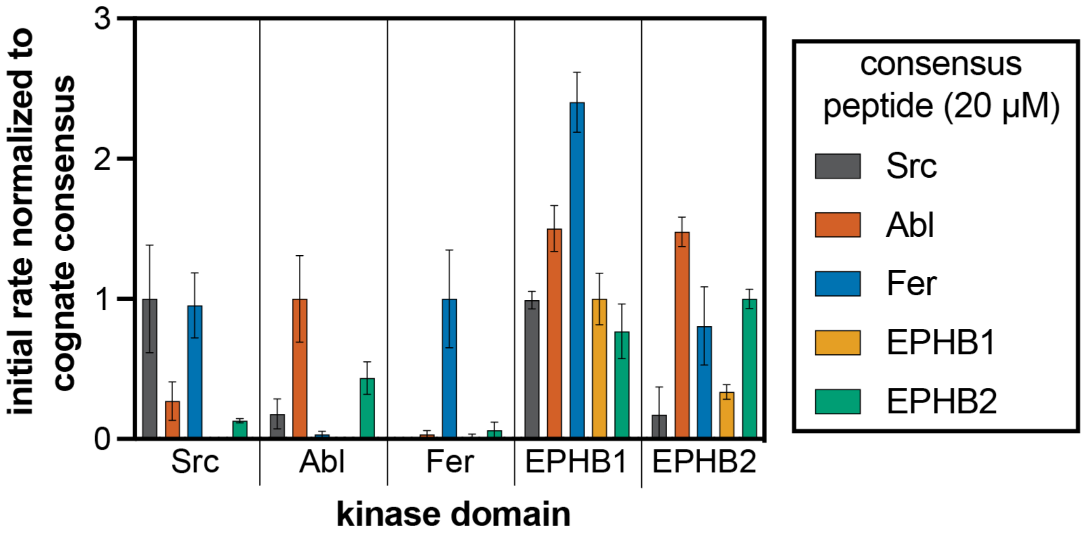


**Figure 2-figure supplement 3. Phosphorylation kinetics of five consensus peptides against five kinases.** Initial rates measured for each kinase were normalized to the rate of the corresponding consensus peptide. All peptides were used at a concentration of 20 μM, and the kinases were used at a concentration of 10-50 nM. Error bars represent the standard deviation from at least three measurements.

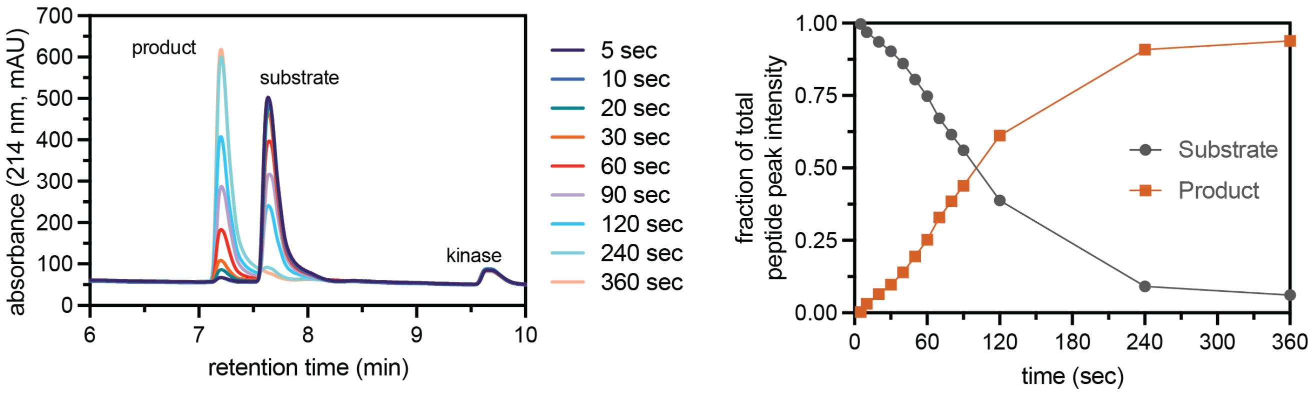


**Figure 3-figure supplement 1. Assay to measure peptide phosphorylation rates using reverse-phase HPLC.** Phosphorylation of the CDK5_Y15 peptide (100 µM) by c-Src (500 nM), monitored by RP-HPLC of selected time points. The HPLC chromatogram shows the formation of a phosphorylated species over time, with concomitant loss of the unphosphorylated peptide (*left*). The area under the two peaks in the chromatogram were quantified and plotted for each time point (*right*). Initial phosphorylation rates were extracted by fitting a line to the first few timepoints.

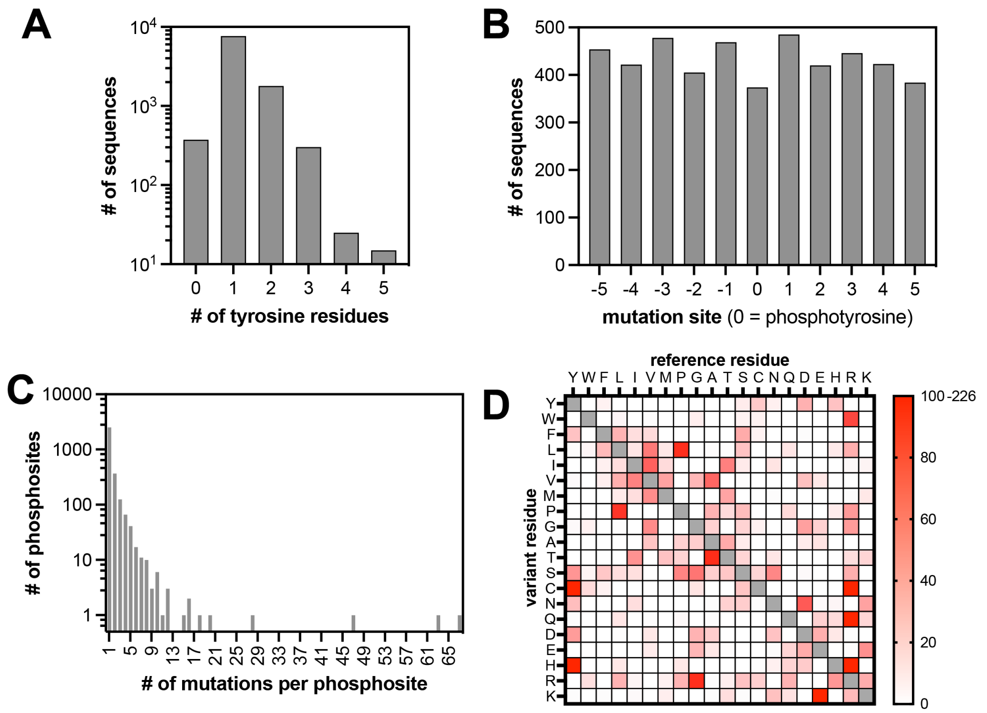


**Figure 4-figure supplement 1. Properties of the pTyr-Var library.** (**A**) Frequency of sequences in the library with different numbers of tyrosine residues. (**B**) Positions of mutations across the library relative to the central tyrosine (zero-position). (**C**) Frequency of substitutions associated with each phosphosite. (**D**) Abundance of each possible amino acid substitution across the library.

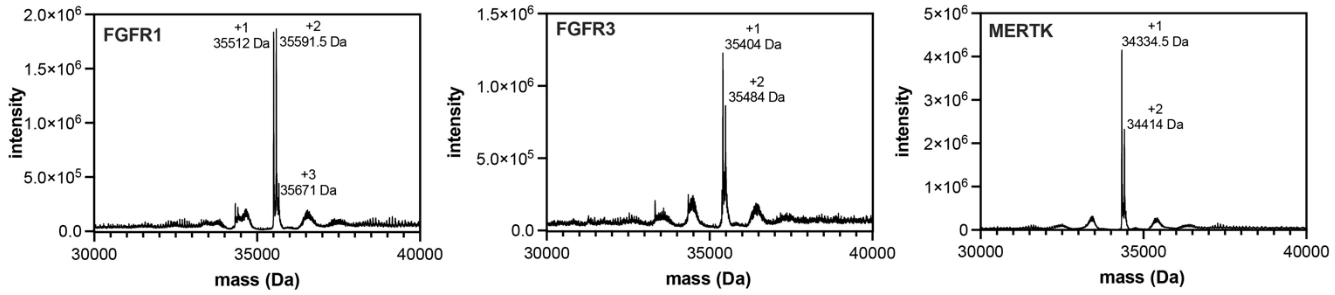


**Figure 4-figure supplement 2. Pre-activation of FGFR1, FGFR3, and MERTK by auto-phosphorylation.** Kinases (25 µM) were incubated with ATP (5 mM) in a magnesium-containing neutral pH buffer for 0.5 to 2 hours, then desalted and concentrated to remove excess ATP. Proteins were analyzed by electrospray-ionization mass spectrometry. The envelope of multiply-charged states was deconvoluted using the instrument software, and the deconvoluted spectra are shown. The number of phosphorylation events on each kinase is labeled.

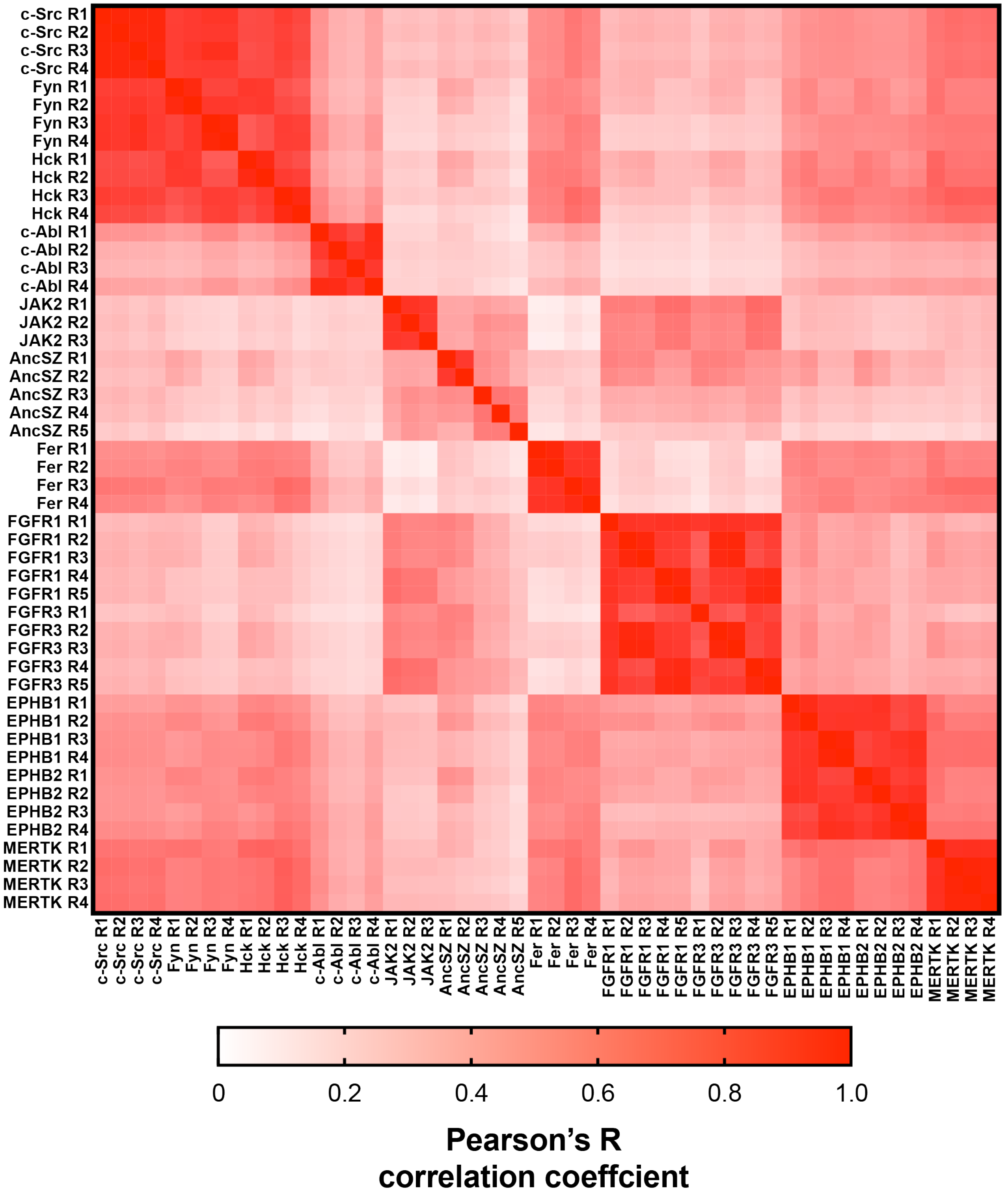


**Figure 4-figure supplement 3. Matrix of Pearson’s correlation coefficients for all replicates of pTyr-Var screens across all 12 kinases.**

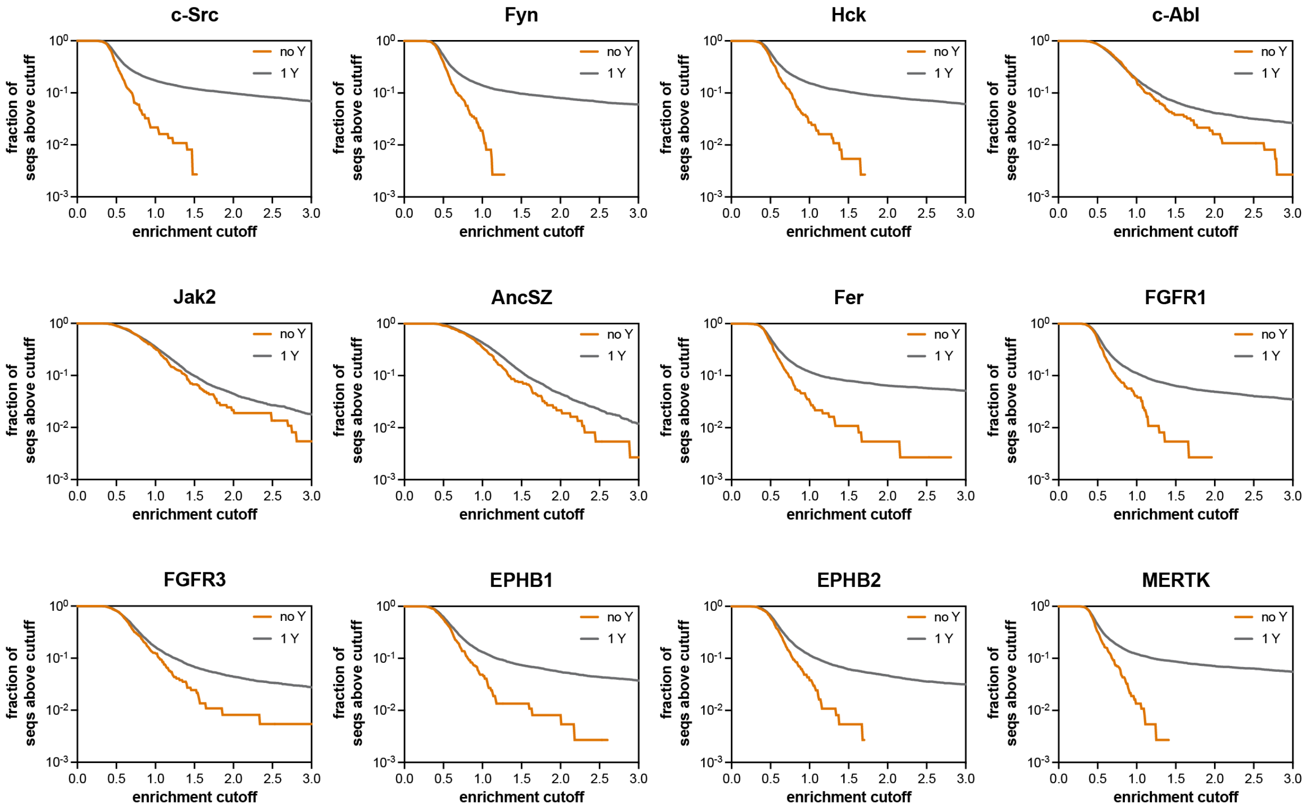


**Figure 4-figure supplement 4. Assessment of the extent of enrichment in pTyr-Var screens with 12 kinases.** These graphs assess what fraction of the sequences containing no Tyr residue (out of 370 sequences) and what fraction of the sequences containing 1 Tyr residue (out of 7,468 sequences) have an enrichment score above the cutoff value indicated on the x-axis.

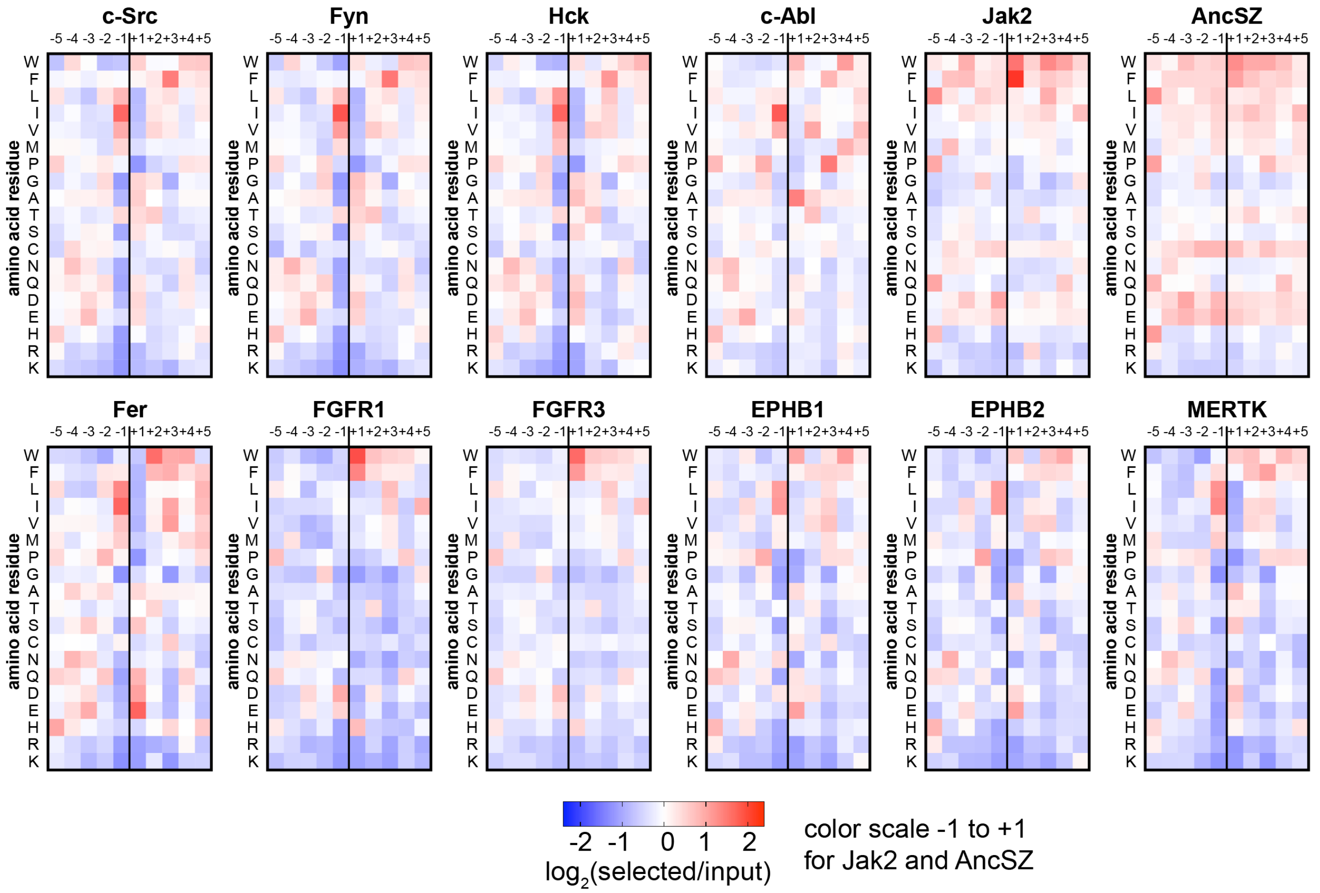


**Figure 4-figure supplement 5. Heatmaps depicting the position-specific amino acid preferences for 12 tyrosine kinase domains, extracted from screens with the pTyr-Var library.** Only sequences with a single central tyrosine were considered in this analysis. Position-specific amino acid enrichment scores were calculated by taking the average $\log_2$-transformed enrichment of every sequence with that particular feature. Values are displayed on a color scale from blue (disfavored sequence features, negative value), to white (neutral sequence features, near zero value), to red (favored sequence features, positive value). Values in the heatmaps are the average of three to five replicates.

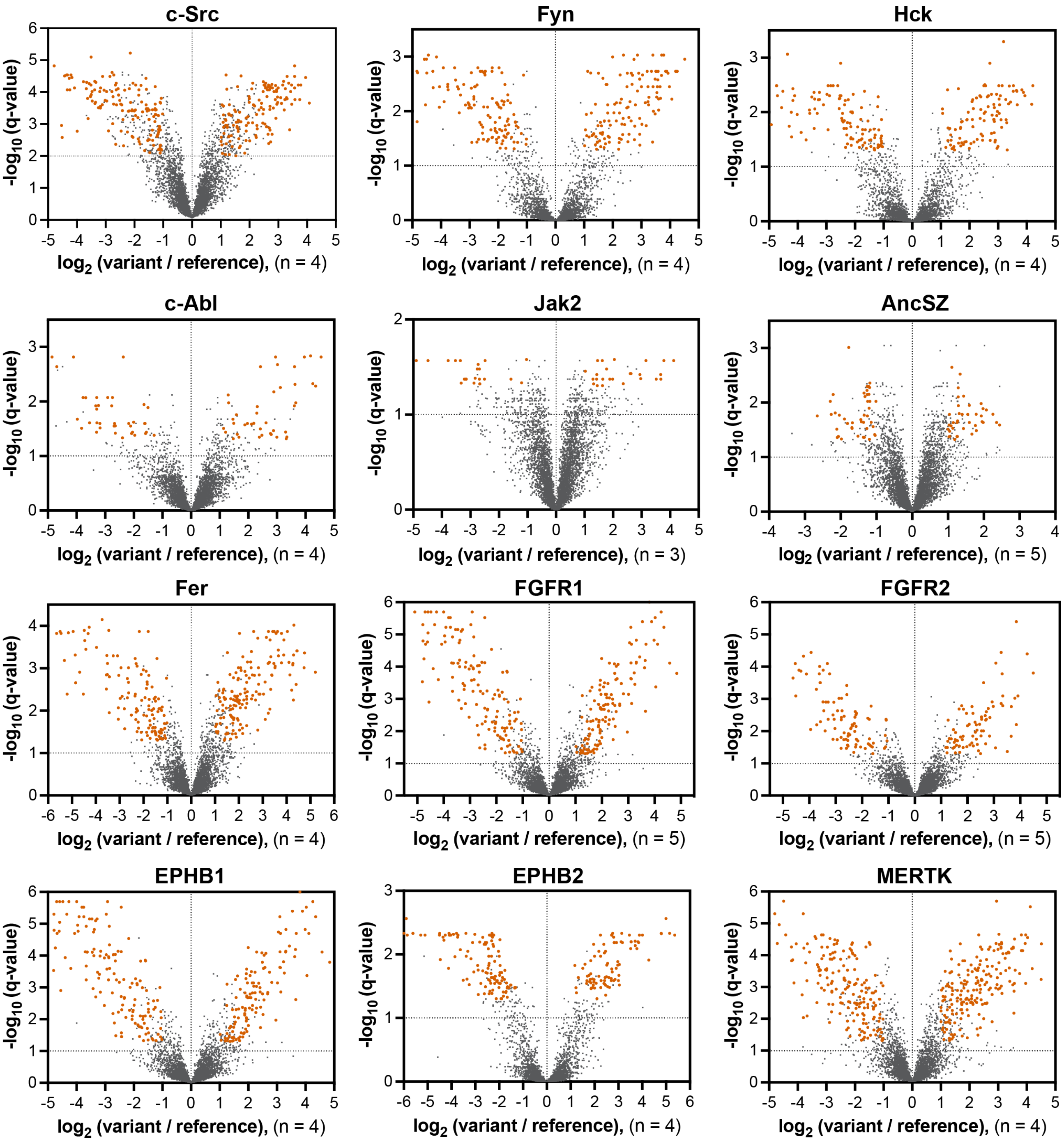


**Figure 4-figure supplement 6. Volcano plots depicting mutational effects in the pTyr-Var screen for 12 kinase domains.** Datasets are the average of three to five replicates. Significant hits are colored in orange-red.

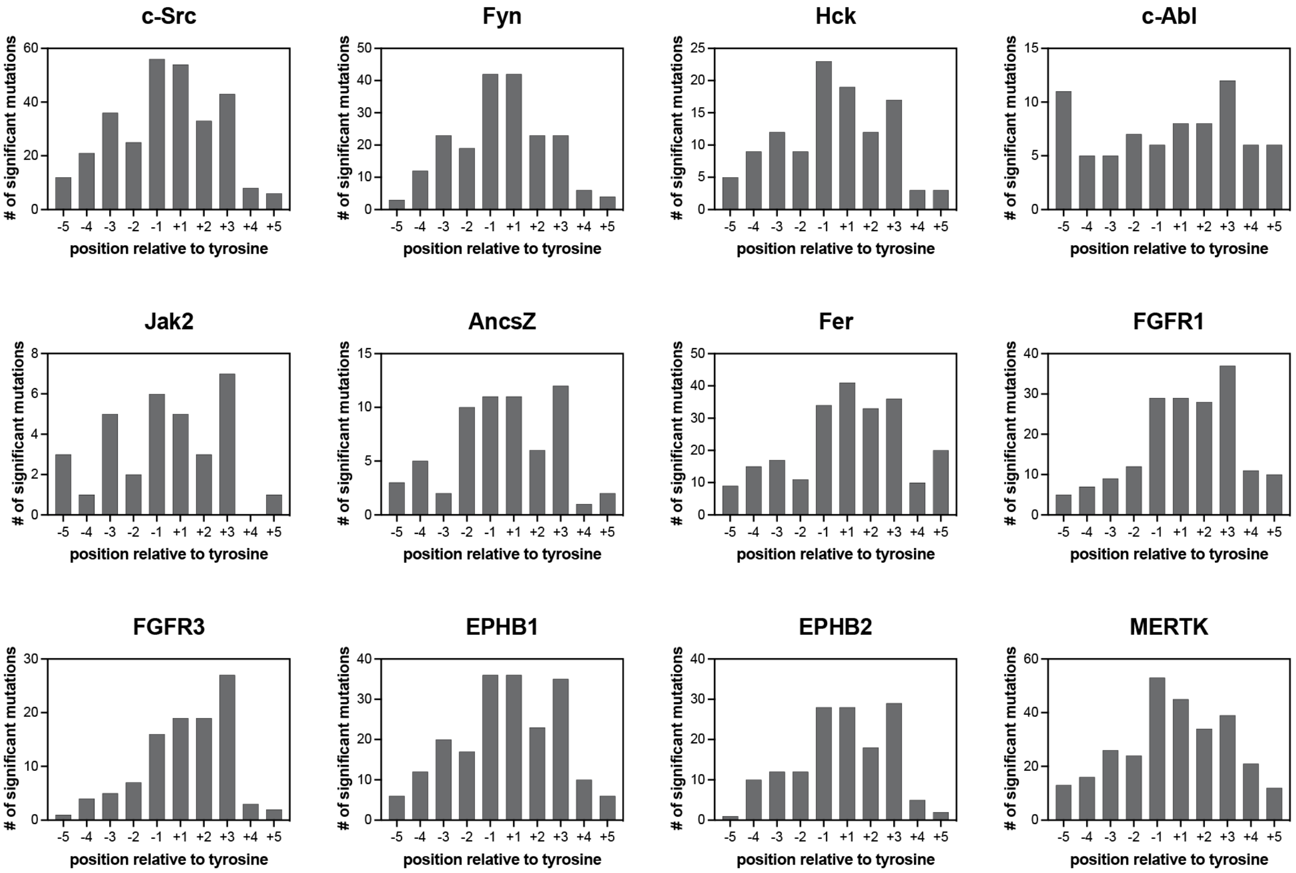


**Figure 4-figure supplement 7. Number of significant mutations for each kinase at each position surrounding the central tyrosine residue.** Mutations that added or removed a tyrosine residue were excluded from these counts.

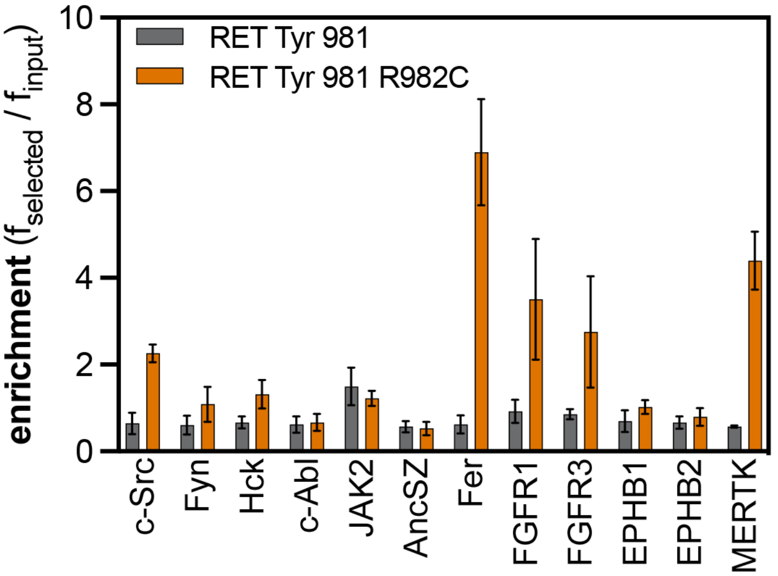


**Figure 4-figure supplement 8. Enrichment scores from pTyr-Var screens for phosphorylation of the RET Tyr 981 reference and variant (R982C) peptides by 12 tyrosine kinases.** Error bars represent the standard deviations from three to five replicates.

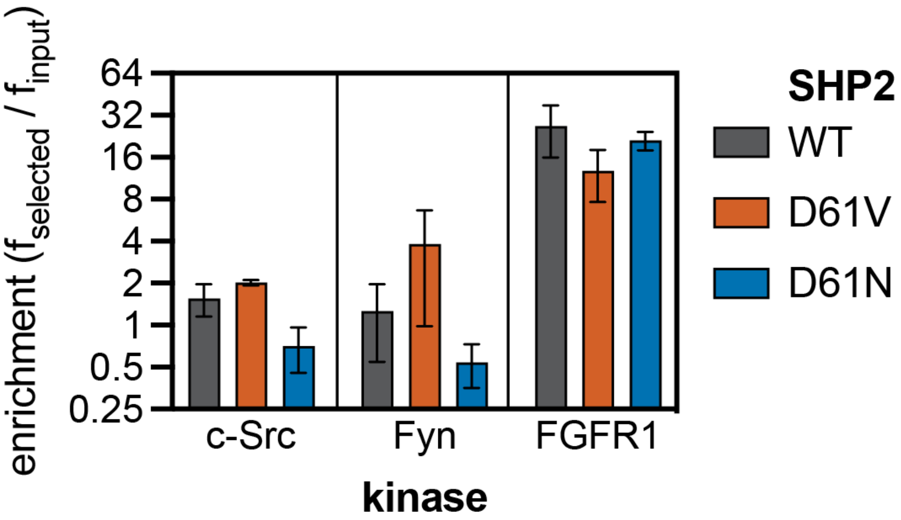


**Figure 4-figure supplement 9. Enrichment scores from pTyr-Var screens for phosphorylation of SHP2 Y62 reference and variant (D61N and D61V) peptides by c-Src, Fyn, and FGFR1.** Error bars represent the standard deviations from three to five replicates.

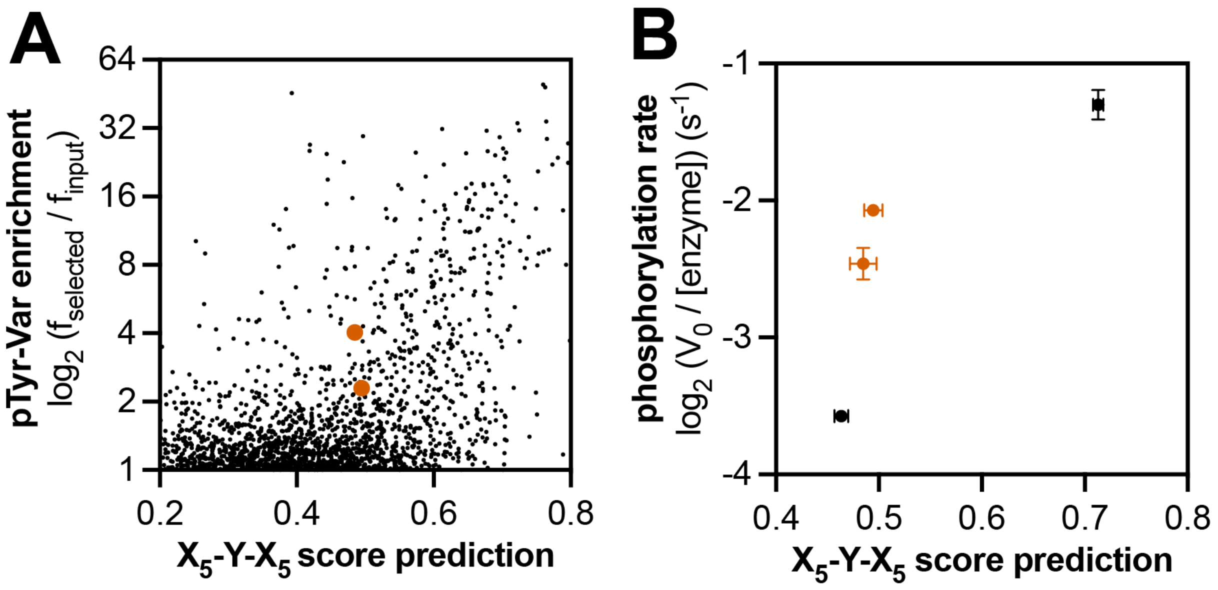


**Figure 5-figure supplement 1. Context-dependent effects of c-Abl substrate recognition. (A)** Correlation of enrichment scores measured for c-Abl in the pTyr-Var library screen with scores predicted from the $X_5$-Y-$X_5$ library using a position-specific scoring matrix. **(B)** Correlation between predicted scores and measured phosphorylation rates for 4 peptides (100 µM) with c-Abl (500 nM). Peptides that showed significant enrichment in the pTyr-Var screen but lower than expected scores from the $X_5$-Y-$X_5$ data are highlighted in orange.

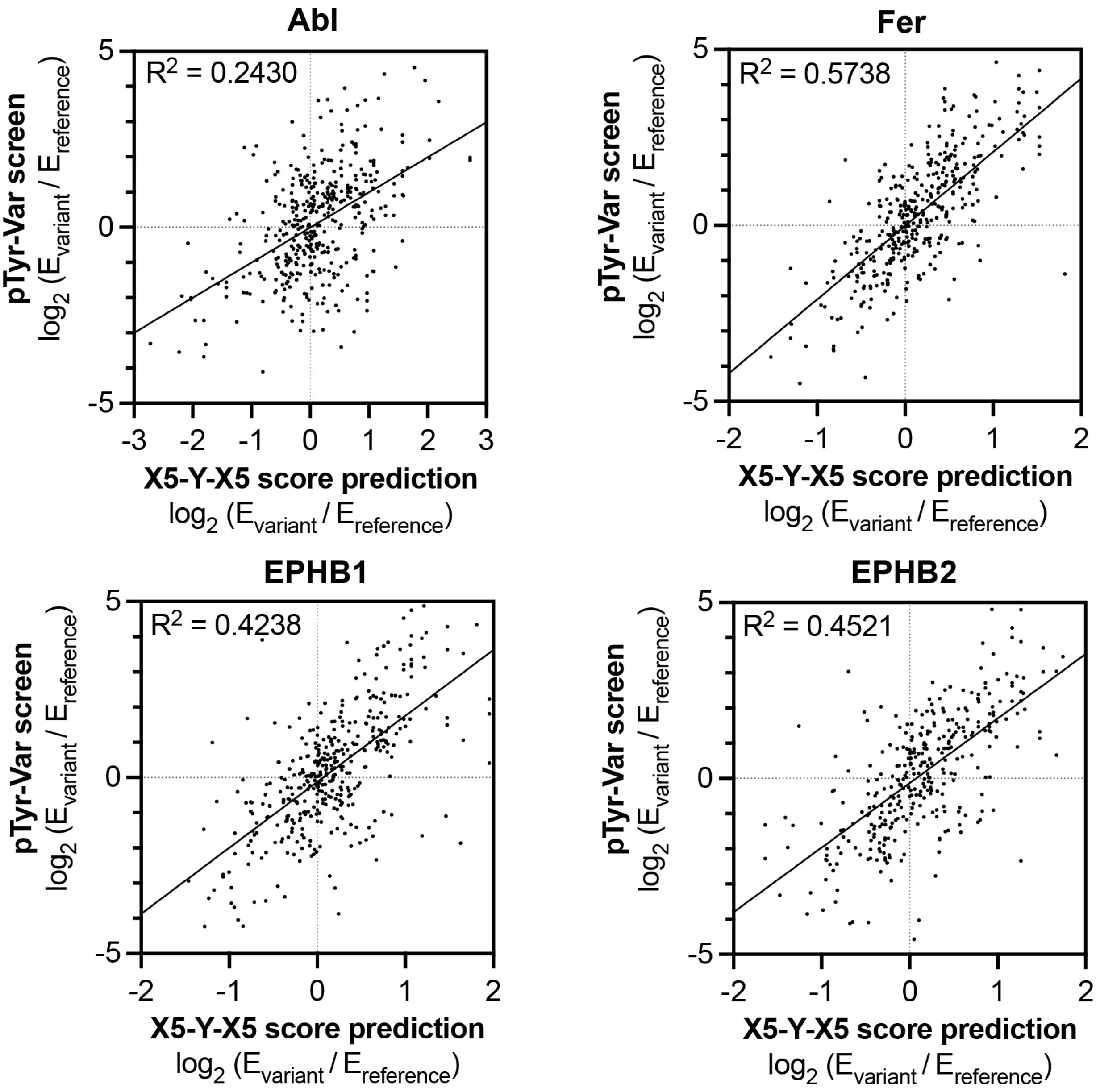


**Figure 5-figure supplement 2. Correlation of variant effects measured in the pTyr-Var library screen with those predicted from the $X_5$-Y-$X_5$ library screen for c-Abl, Fer, EPHB1, and EPHB2.** Several points lie in the top-left and bottom-right quadrant, indicating a discrepancy between the measured mutational effect in the pTyr-Var screen and the predicted mutational effect from the $X_5$-Y-$X_5$ screen.

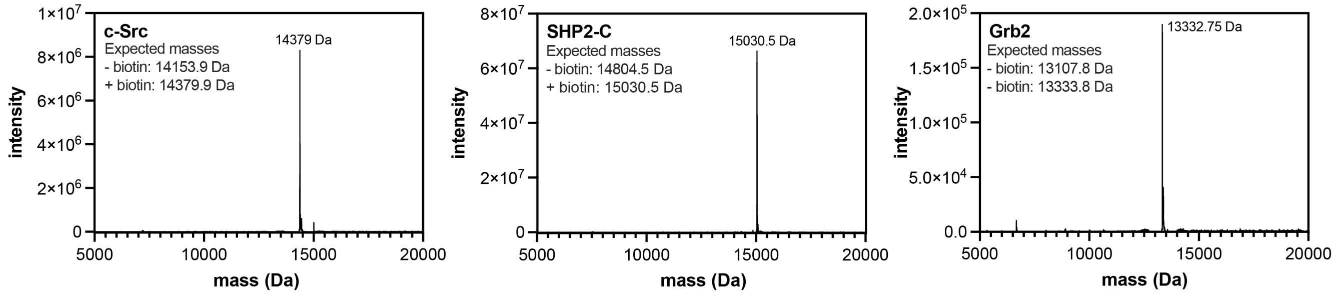


**Figure 6-figure supplement 1. Mass spectrometry analysis of biotinylated SH2 domains.** Proteins were analyzed by electrospray-ionization mass spectrometry. The envelope of multiply-charged states was deconvoluted using the instrument software, and the deconvoluted spectra are shown.

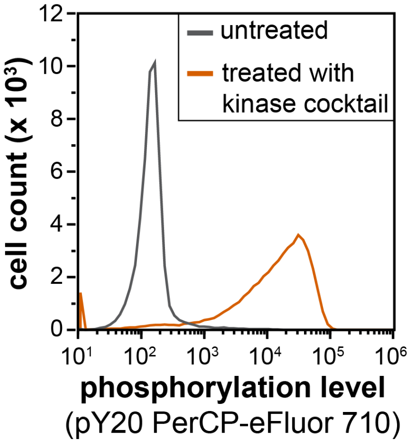


**Figure 6-figure supplement 2. Flow cytometry analysis of library phosphorylation by a cocktail of tyrosine kinases.** Cells displaying the $X_5$-Y-$X_5$ library were treated with a kinase cocktail containing c-Src, c-Abl, AncSZ, and EPHB1 for 3 hours, then labeled with a pan-phosphotyrosine antibody (PY20 PerCP-eFluor 710) and analyzed by flow cytometry.

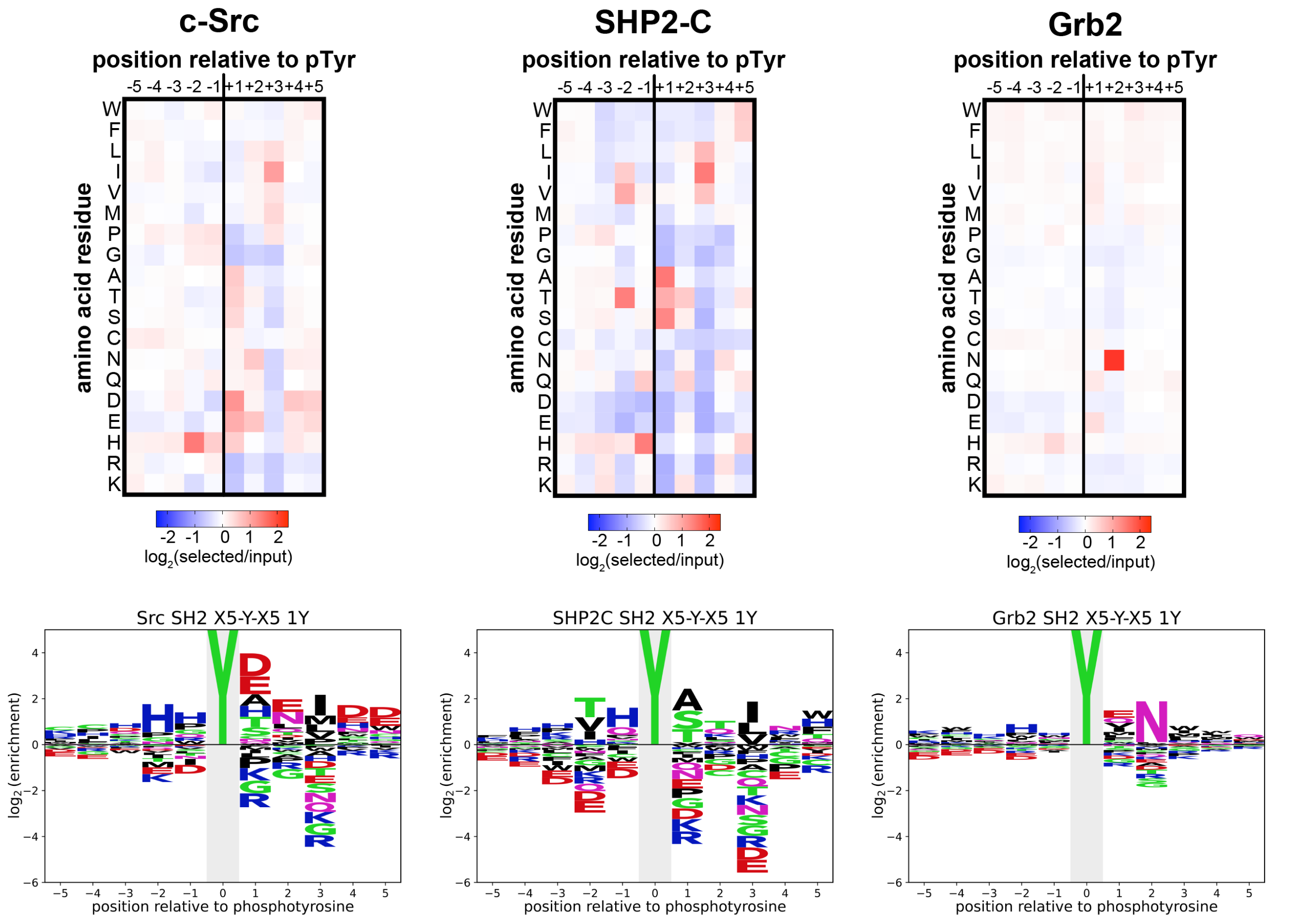


**Figure 6-figure supplement 3. Heatmaps and logos depicting the specificities of the c-Src, SHP2-C, and Grb2 SH2 domains, measured using the $X_5$-Y-$X_5$ library.** Only peptides with one central tyrosine were considered in this analysis. Enrichment scores were $\log_2$-transformed and are displayed on a color scale from blue (disfavored sequence features, negative value), to white (neutral sequence features, near zero value), to red (favored sequence features, positive value). The same values were used to plot the heatmaps and the sequence logos. The height for the central "Y" in the sequence logos is an arbitrary value, chosen for optimal visualization of other features. Values are the average of three replicates.

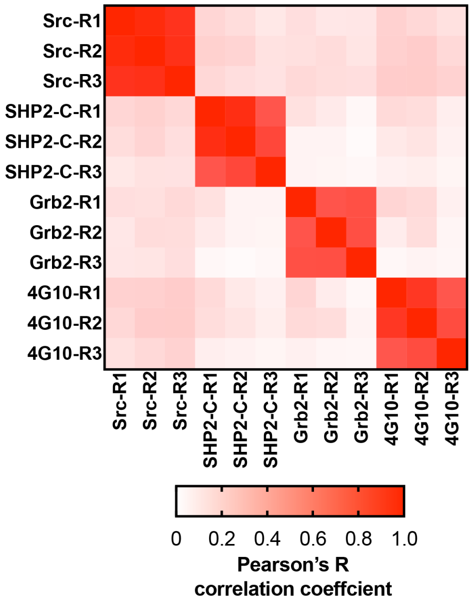


**Figure 6-figure supplement 4. Matrix of Pearson's correlation coefficients for all replicates of pTyr-Var screens across all 3 SH2 domains and 4G10 platinum.**

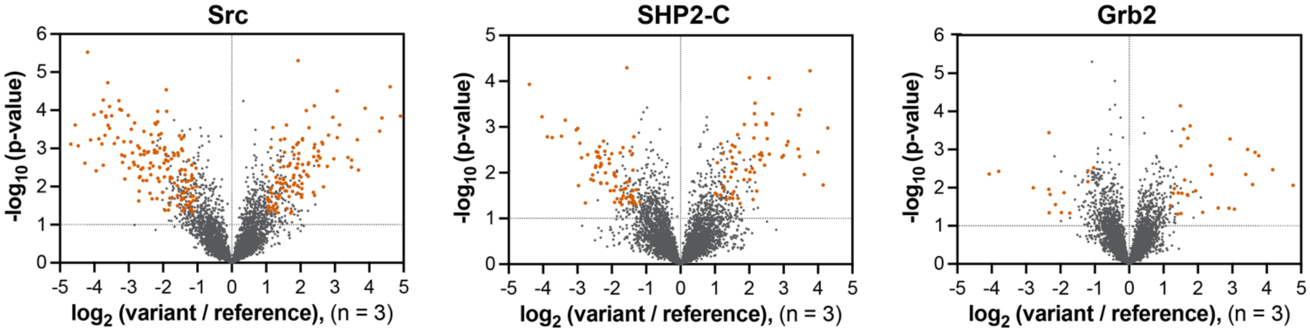


**Figure 6-figure supplement 5. Volcano plots depicting mutational effects in the pTyr-Var screen for 3 SH2 domains.** Datasets are the average of three replicates. Hits are colored in orange-red.

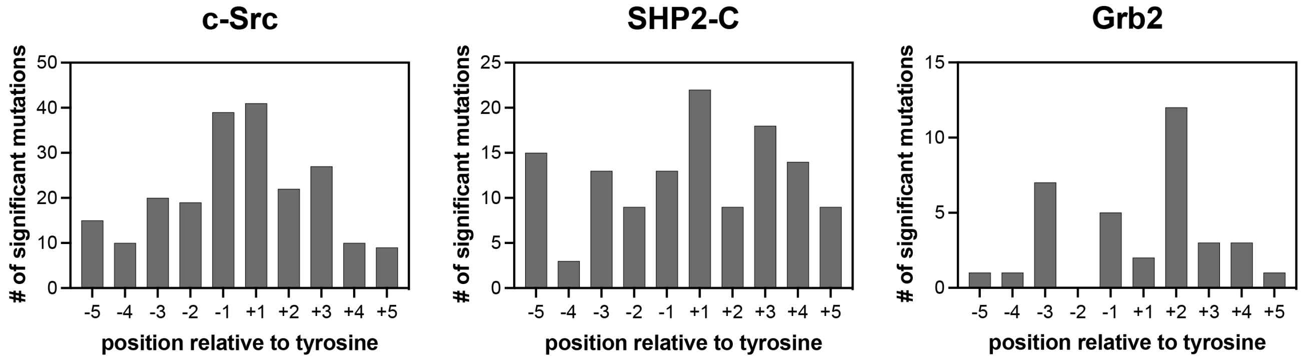


**Figure 6-figure supplement 6. Number of significant mutations for each SH2 domain at each position surrounding the central phosphotyrosine residue.** Mutations that added or removed a tyrosine residue are excluded from these counts.

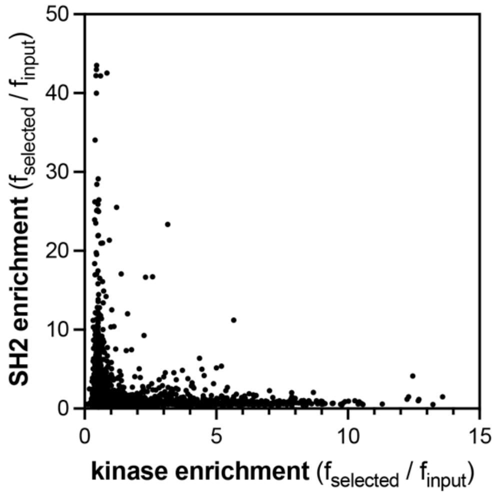


**Figure 6-figure supplement 7. Comparison of the pTyr-Var screens for the c-Src kinase and SH2 domains.** Kinase domain data are the average of four replicates, and SH2 data are the average of three replicates.

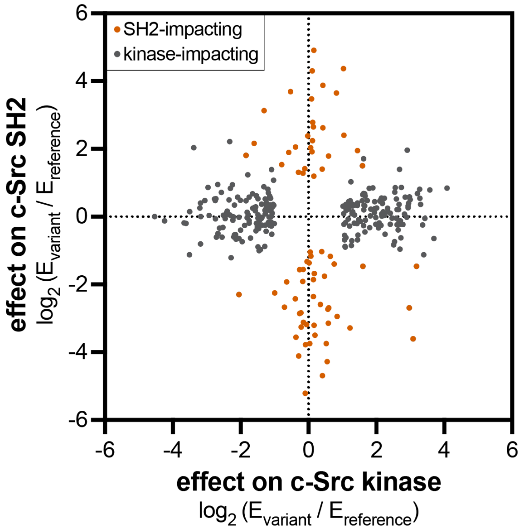


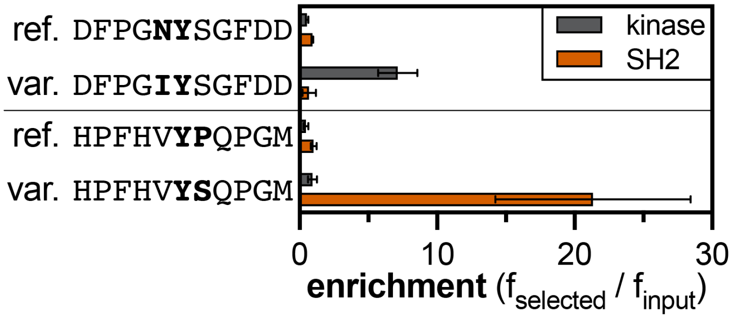


**Figure 6-figure supplement 8. Divergent effects of phosphosite-proximal mutations on c-Src kinase and SH2 domain recognition.** The graph on the left shows the effects of mutations that were significant for the kinase domain (gray) or the SH2 domain (orange-red). The graph on the right shows examples of phosphosite-proximal mutations selectively impact the kinase or SH2 domain of c-Src. Error bars for the kinase and SH2 domain indicate the standard deviations from four and three replicates, respectively.

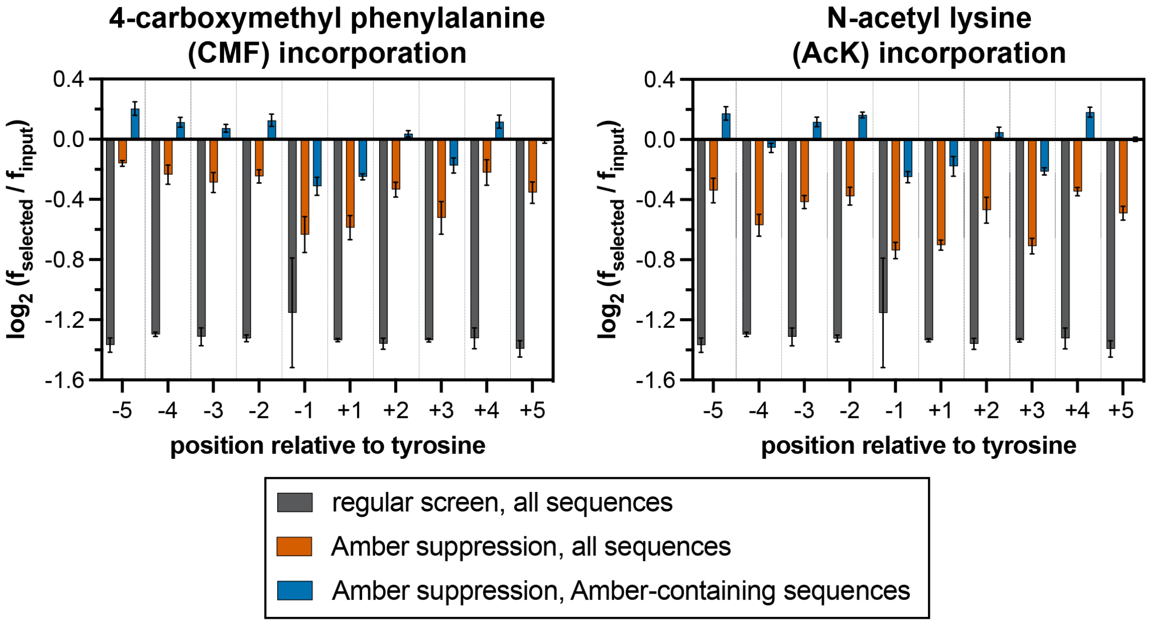


**Figure 7-figure supplement 1. Stop codon enrichment levels in c-Src $X_5$-Y-$X_5$ screens using different analysis methods.** Error bars represent the standard deviations from three screens.

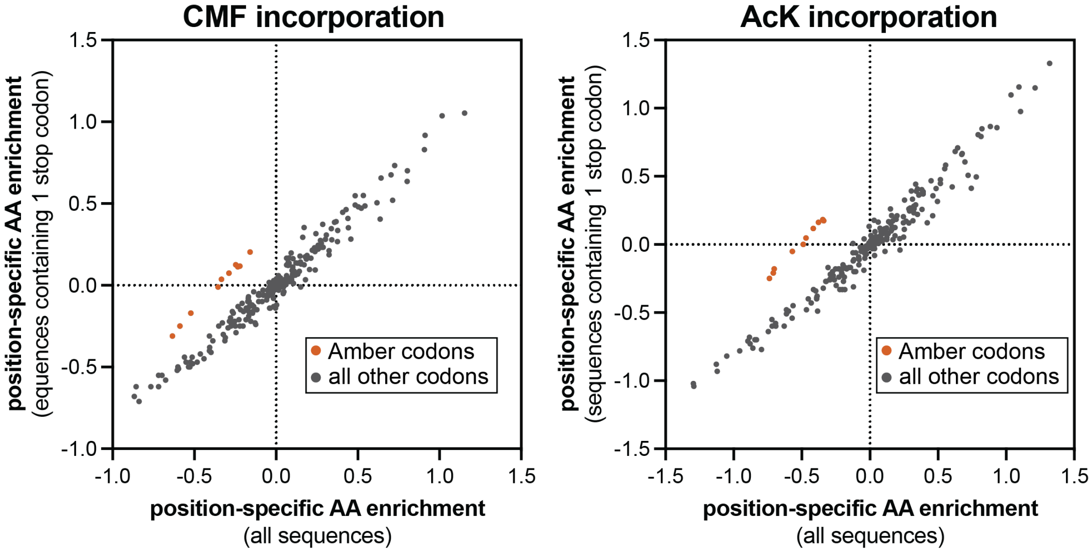


**Figure 7-figure supplement 2. Comparison of position-specific enrichments in screens with Amber suppression analyzed in two different ways.** In each plot, the enrichment of specific amino acids or a stop codon, after phosphorylation by c-Src and bead-based selection, were calculated using two different methods. X-values indicate log-transformed enrichment values calculated across all sequences in the library. Y-values indicate log-transformed enrichment values only for sequences that contain exactly one Amber stop codon. The orange-red points correspond to the Amber codon enrichments at all 10 positions, which selectively fall off of the x = y diagonal line.

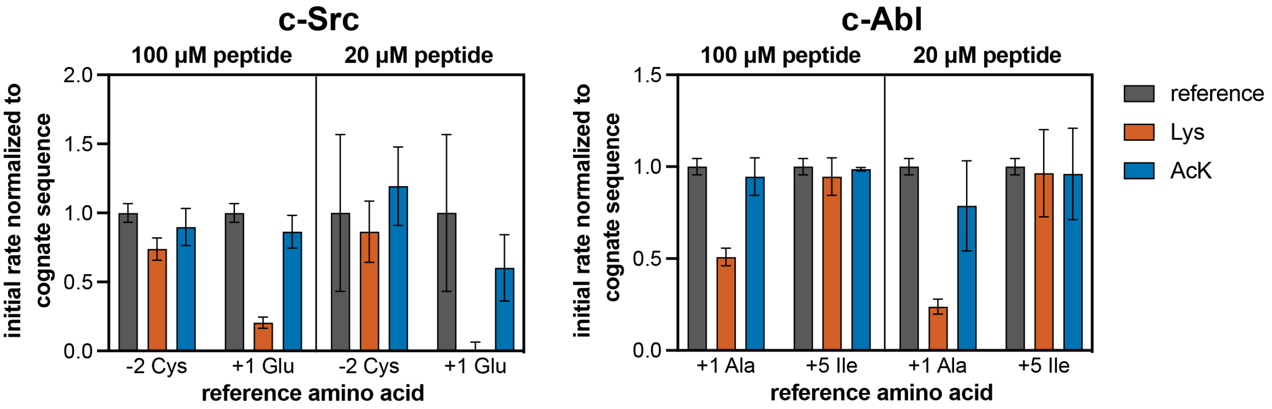


**Figure 7-figure supplement 3. Phosphorylation kinetics of Lys- and AcK-containing consensus peptides against c-Src and c-Abl.** Initial rates measured for each kinase were normalized to the rate of the corresponding cognate consensus peptide. Peptides were used at a concentration of 100 or 20 µM, and the kinases were used at a concentration of 10-50 nM. Error bars represent the standard deviation from three measurements.
